# Involuntary shifts of spatial attention contribute to distraction - Evidence from oscillatory alpha power and reaction time data

**DOI:** 10.1101/2020.06.29.161992

**Authors:** Annekathrin Weise, Thomas Hartmann, Fabrice Parmentier, Nathan Weisz, Philipp Ruhnau

## Abstract

Imagine you are focusing on the traffic on a busy street to ride your bike safely when suddenly you hear the siren of an ambulance. This unexpected sound involuntarily captures your attention and interferes with ongoing performance. We tested whether this type of distraction involves a spatial shift of attention. We measured behavioral data and magnetoencephalographic alpha power during a crossmodal paradigm that combined an exogenous cueing task and a distraction task. In each trial, a task-irrelevant sound preceded a visual target (left or right). The sound was usually the same animal sound (i.e., standard sound). Rarely, it was replaced by an unexpected environmental sound (i.e., deviant sound). Fifty percent of the deviants occurred on the same side as the target, and 50% occurred on the opposite side. Participants responded to the location of the target. As expected, responses were slower to targets that followed a deviant compared to a standard. Crucially, this distraction effect was mitigated by the spatial relationship between the targets and the deviants: responses were faster when targets followed deviants on the same versus different side, indexing a *spatial* shift of attention. This was further corroborated by a posterior alpha power modulation that was higher in the hemisphere ipsilateral (vs. contralateral) to the location of the attention-capturing deviant. We suggest that this alpha power lateralization reflects a spatial attention bias. Overall, our data support the contention that spatial shifts of attention contribute to deviant distraction.

## 1. Introduction

Efficient cognitive functioning requires the ability to filter out task-irrelevant events to effectively perform a task at hand. On the other hand, it may also be adaptive to detect unexpected but behaviorally relevant changes in the soundscape. Hence, selective attention and change detection must be carefully balanced to limit the negative impact of task-irrelevant stimuli while allowing for potentially important stimuli to break through attentional filters. One upshot of this balance is the cognitive system’s vulnerability to distraction. For example, task-irrelevant events, such as an ambulance siren while you are riding your bike, may distract you and affect driving performance.

In an experimental setting, unexpected task-irrelevant events, so-called deviants, usually violate sensory predictions by deviating from an otherwise (more or less) regular sound sequence consisting of so-called standards. Distraction due to auditory deviants has been extensively studied via event-related potentials and behavioral measures (e.g., Bendixen et al., 2007; Schröger & Wolff, 1998; Wetzel et al., 2012). It is well known that deviant distraction is accompanied by distinct event-related markers (for reviews, see Escera et al., 2000; Parmentier, 2014) and a behavioral effect observed when unexpected sounds violate sensory predictions (e.g., Bendixen et al., 2008; Parmentier et al., 2011). Distraction on behavioral level is defined as the slowing of responses during the primary task following the presentation of a task-distracting event compared to standard events. This effect is thought to result from an involuntary (stimulus-driven, bottom-up, or exogenous) shift of attention from the task towards the deviant and back to the task, which results in a time penalty (Parmentier et al., 2008; Schröger, 1996) and appears to be related to a transient inhibition of action (Dutra et al., 2018; Vasilev et al., 2019; Wessel and Aron, 2017).

The exact nature of the attention shift in crossmodal scenarios, in which the auditory deviant and the visual target occur at different locations, still needs to be established. One hypothesis proposed is that of a spatial shift (Parmentier et al., 2008; Parmentier, 2014). According to that hypothesis, deviant sounds may capture attention and trigger an involuntary shift of attention to the location of the deviant sound, and from there to the target’s location upon its appearance. Because spatial shifts require time to complete (Cheal & Gregory, 1997; Luck et al., 1996; Shiu & Pashler, 1994), they might contribute to distraction by deviant sounds (Parmentier, 2014).

Support for the spatial shift hypothesis is currently lacking. The few reports relevant to this issue cannot rule out alternative (though not mutually exclusive) interpretations (Parmentier et al., 2008), or present inconsistent results (Corral & Escera, 2008). However, indirect support can be found in behavioral data outside of the deviant distraction literature. This is for example the case in studies that applied an exogenous spatial cueing paradigm in which a task-irrelevant sound, such as a noise burst, appeared lateralized (left or right) briefly before the presentation of an ipsi- or contralateral visual target. While the location of the sound does not predict the location or the type of upcoming visual target, results show a performance advantage when cue and visual target appeared on the same side (Dufour, 1999; Feng et al., 2014; McDonald et al., 2000; Spence & Driver, 1997). This pattern of results indicates that cues are effective in generating an involuntary shift of spatial attention.

Some of the studies showing behavioral evidence for an involuntary shift of spatial attention draw further support from electroencephalographic (EEG) data (Feng et al., 2017; Störmer et al., 2016). Focusing on oscillatory activity, the authors found a sound-induced modulation in posterior alpha power that built up rapidly and was rather short-lived, thus resembling the properties of involuntary attention (Corbetta & Shulman, 2002). Crucially, the posterior alpha power modulation was lateralized. That is, the alpha power is higher in the hemisphere ipsilateral to the location of the attention-capturing sound (compared to contralateral alpha power). In fact, the posterior alpha power lateralization is a well-known finding from research on voluntary (goal-directed, top-down, or endogenous) attention that has been linked to spatial attention bias (e.g., Banerjee et al., 2011; Deng et al., 2019; Rihs et al., 2009; Sauseng et al., 2005; Thut et al., 2006; Worden et al., 2000; Wöstmann et al., 2016). Research on voluntary attention typically uses endogenous spatial cueing tasks, in which cues require participants’ to shift their attention to certain target locations (left or right). Irrespective of whether spatial attention is voluntarily directed to a visual or auditory target, sustained oscillatory alpha power appears to be lateralized, especially over parieto-occipital areas. This lateralization is often driven by an ipsilateral increase and/or a contralateral decrease in alpha power relative to the attended side, thought to reflect the inhibition of brain regions that process task-irrelevant information at unattended locations, or facilitate processing of task-relevant information at attended locations, respectively (for reviews, see Hanslmayr, 2011; Klimesch, 2012; Peylo et al., in press).

### Current approach

To test the possible role of a spatial shift of attention in deviance distraction, we employed a crossmodal paradigm combining auditory distraction and an exogenous spatial cueing task (Figure 1). In each trial, a task-irrelevant sound preceded a visual target. In most trials (80%), the same standard sound was used. In the remaining trials (20%), deviant sounds were used. In the Deviant Left condition, deviants were presented to the participant’s left ear. In the Deviant Right condition, they were presented to the right ear. As in an exogenous spatial cueing task, the location of the deviant was non-informative about the spatial location of the upcoming target and could either be congruent or incongruent with it (with equal probabilities). That is, 50% of the deviants were presented on the same side as the upcoming visual target (Congruent condition), while the remaining 50% of the deviants were presented on the side opposite to the upcoming visual target (Incongruent condition). Participants were instructed to respond to the target location (left vs right).

**Figure 1:**
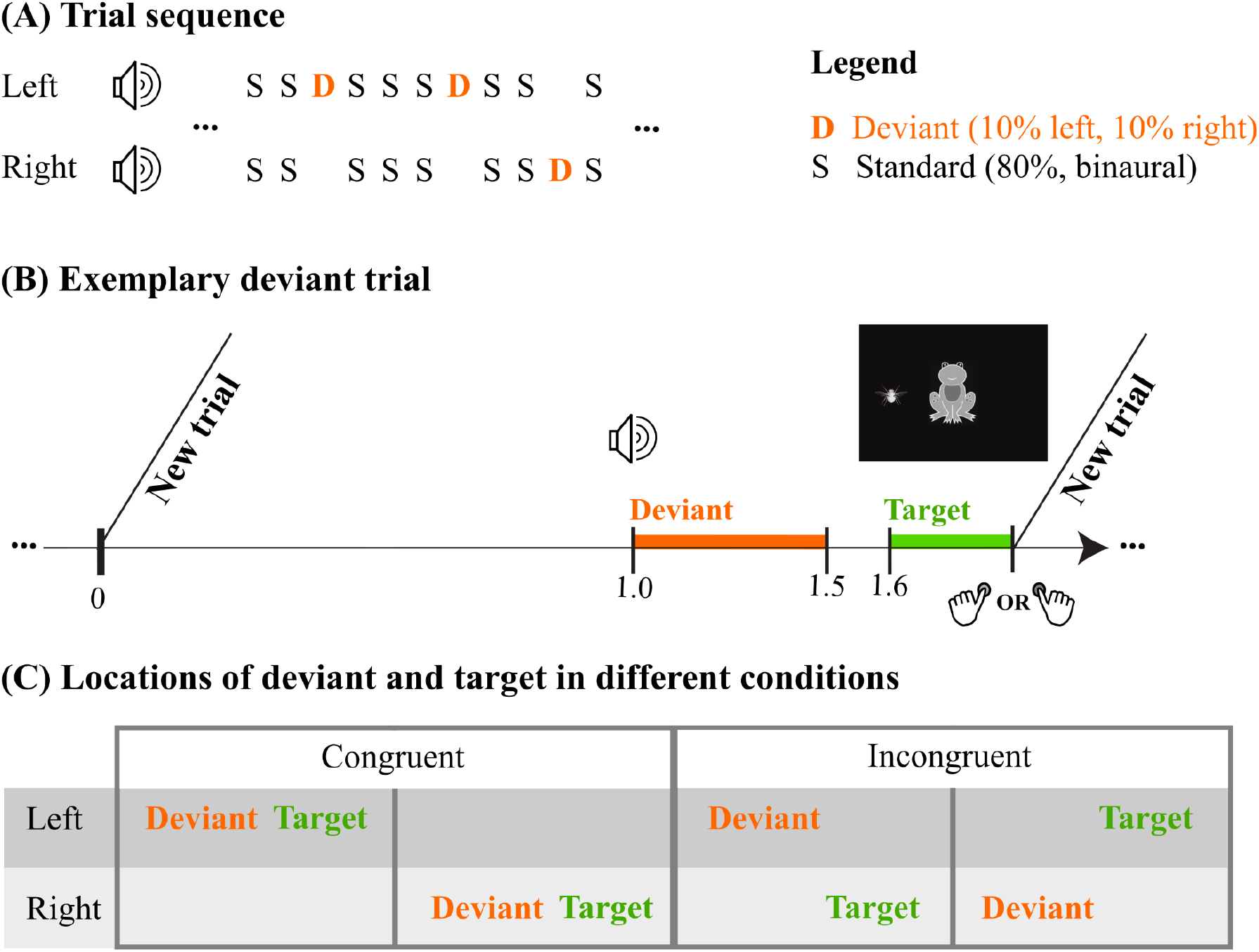
Cross-modal paradigm combining a deviant distraction task with an exogenous spatial cueing task. (A) Exemplary trial sequence. Trials could either be standard trials or deviant trials depending on whether the task-irrelevant sound was a binaurally presented standard sound (p = 0.8) or a lateralized deviant sound presented either left (p = 0.1) or right (p = 0.1). (B) Exemplary deviant trial. During each trial, the task-irrelevant sound - here a deviant sound, illustrated as an orange bar - preceded the task-relevant visual stimulus, illustrated as a green bar. Participants had to indicate whether the target occurred to the left or right of the fixation frog. (C) Visual representation of the two spatial conditions (Congruent and Incongruent) and the corresponding spatial locations (Left, Right) of deviants and targets.

At the behavioral level, our predictions were two-fold. First, we expected to find longer reaction times (RTs) to visual targets following a deviant compared to a standard, reflecting an involuntary shift to and from the deviant (Escera et al., 1998; Parmentier et al., 2008; Parmentier, 2014; Ruhnau et al., 2013). Secondly, and more crucially, we predicted a *spatial* congruence effect whereby RTs would be shorter for congruent than incongruent trials.

To examine brain activity, we recorded MEG data to shed light on the neural oscillatory activity at play in our task, with an emphasis on alpha power and its link to involuntary spatial attention. We contrasted the conditions Deviant Left minus Deviant Right. We predicted that the deviant would induce a lateralized, short-lived, alpha power modulation in parieto-occipital areas that would build up rapidly after stimulus onset, thereby matching the characteristics of involuntary spatial attention (Feng et al., 2017; Störmer et al., 2016). More specifically, with respect to the alpha power modulation, we expected to find relatively higher alpha power in the hemisphere ipsilateral to the attention-capturing deviant sound and relatively lower alpha power in the hemisphere contralateral to the attention-capturing deviant. These hypotheses are in agreement with the literature on spatial attention (involuntary spatial attention: Feng et al., 2017; Störmer et al., 2016; voluntary spatial attention: e.g. Wöstmann et al., 2019).

## 2. Methods

The study was approved by the local ethics committee of the Paris Lodron Universität Salzburg and adhered to the code of ethics of the World Medical Association (Declaration of Helsinki).

### 2.1 Participants

We analyzed the data from 30 healthy young adults who participated in the MEG experiment (12 males, mean age ± STD: 27 ± 6 yrs, 4 left-handed, 1 ambidextrous). Data of one additional participant were recorded but excluded from analysis due to excessive artifacts in the MEG. Volunteers gave written, informed consent prior to their participation. All participants reported normal hearing and normal or corrected-to-normal vision, and none reported a history of neurological or psychiatric diseases. Volunteers received 10€/hour (h) or 1 credit point/h as compensation for their participation.

### 2.2 Stimuli and Procedure

We used auditory stimuli and visual stimuli. Auditory stimuli were either standards (p = .8) or deviant (p = .2) sounds. Both had a duration of 0.5 s (including 10 ms rise and fall times). The standard sound was that of a buzzing mosquito. Standards were binaurally presented. The deviant sound was chosen randomly for each participant from a pool of 56 different environmental sounds (e.g., speech, animal voices, tool noises, etc.; selected from a commercial CD: 1,111 Geräusche, Döbeler Cooperations, Hamburg, Germany). Deviants were monaurally presented. All sounds were equalized for overall intensity (RMS). The auditory stimulation was delivered via MEG-compatible pneumatic in-ear headphones (SOUNDPixx, VPixx technologies, Canada). The auditory stimuli were presented at 50 dB above the sensation threshold level. Each participant’s individual sensation threshold was determined using a 200-ms 1000 Hertz (Hz) tone presented via a bayesian threshold method (Kontsevich & Tyler, 1999; Sanchez et al., 2016) utilizing the VBA toolbox (Daunizeau et al., 2014). The sensation threshold was measured separately for the right and left ears before the start of the experiment.

Visual stimuli consisted in cliparts of a frog and of a fly. The frog was presented in the center of the screen and served as a fixation point. The size of the frog was 2.27° x 2.64°. Its color contained several levels of gray (see Figure 1) and its estimated perceived luminance was 200.91. The fly served as a target. The size of the fly was 1.47° x 1.23°. Its color contained several levels of gray and its luminance was 214.34. In each trial, the fly was presented on the left or on the right of the frog (with equal probabilities). The visual angle between the center of the screen and the target was 8.93°. The visual stimuli were presented inside of the magnetically shielded room using a projector (PROPixx DLP LED Projector), via a back-projection screen and a mirror system.

The experiment was programmed in MATLAB 9.1 (The MathWorks, Natick, Massachusetts, U.S.A) using the open-source Psychophysics Toolbox (Brainard 1997; Kleiner et al. 2007) and o_ptb, an additional class-based abstraction layer on top of the Psychophysics Toolbox (Hartmann and Weisz, 2020; https://gitlab.com/thht/o_ptb). Stimuli were presented with precise timing using the VPixx System (DATAPixx2 display driver, PROPixx DLP LED Projector, TOUCHPixx response box by VPixx Technologies, Canada). The Blackbox2 Toolkit (The Black Box ToolKit Ltd, Sheffield, UK) was used to measure and correct for timing inaccuracies between triggers and the visual and auditory stimulation. Note that the tubes of the MEG-proof sound system caused a time delay between sound and trigger of 16.5 ms that was corrected within the stimulation protocol.

We employed a cross-modal paradigm combining a distraction task with an exogenous spatial cueing task. Participants performed a visual-spatial two-alternative forced-choice task (see Figure 1). Throughout the task, a frog was displayed in the center of the screen that participants were asked to fixate on at all times. In every trial, a visual target was presented that consisted of the shape of a fly featured next to the frog. This visual target appeared on the left or on the right of the frog with equal probability (the order of these trials was random for every participant). Participants were instructed to hold their left and right thumbs over left/right response buttons respectively and to press these buttons to indicate the side where the visual target (fly) had been presented. The mapping between the response buttons and the target location was counterbalanced across participants. The target remained on the screen until a response was recorded or for a period of 1 s (whichever occurred first). A response within this time window terminated the current trial and initiated the next trial.

Apart from the visual target, each trial also involved the presentation of an auditory stimulus, which participants were instructed to ignore. This sound was always presented after a 0.8 s silent interval at the start of each trial, and its onset preceded that of the visual target by 0.6 s. That is, a silent interval of 0.1 s separated the offset of the sound from the onset of the visual target. This timing was selected based on earlier work (Ruhnau et al., 2013) and because short stimulus onset asynchrony is best suited to study distraction (Bendixen et al., 2010). The sound was either a standard or a deviant. The order of the trials was pseudorandomized such that deviant trials occurred randomly with the constraint that each deviant trial was preceded by at least two standard trials.

Two types of deviant trials, equiprobable (p = .1) were defined. In the Congruent condition, the deviant sound and visual target were presented on the same side (left or right, with equal probabilities). In the Incongruent condition, they were on opposite sides (left/right vs. right/left, with equal probabilities). Each of the 56 deviant sounds occurred only once within a block of trials, and a total of four times in the whole experiment - twice on the left (once in the Incongruent and once in the Congruent condition) and twice on the right (once in the Incongruent and once in the Congruent condition).

The experiment consisted of seven blocks in total, each lasting approximately 5 min. and consisting of 160 trials (128 standards, 16 deviants presented on the left, 16 deviants presented on the right). Thus, the time spent on task (excluding breaks between blocks) could be up to approximately 35 min. Thereafter, participants practiced the task with fewer trials to familiarize themselves with the experiment. These were played backward so that participants did not become familiar with those sounds.

### 2.3 MEG recording and preprocessing

Before beginning the MEG recording, five head position indicator (HPI) coils were placed on the left and right mastoid, on the upper right and lower right forehead, and on the left middle forehead. Anatomical landmarks (nasion and left/right preauricular points), the HPI locations, and at least 300 headshape points were sampled using a Polhemus FASTTRAK digitizer. Six pre-gelled and self-adhesive Ag/AgCl electrodes (Ambu Neuroline 720) were attached to record the vertical and horizontal electrooculogram (EOG) and the electrocardiogram (ECG). One additional electrode - placed at the right shoulder - served as a ground.

Participants were in a seated position during data recording. A chin rest was used to avoid head movements during and across different blocks of recording. The eye, heart, and magnetic signals of the brain were recorded at 1000 Hz (hardware filters: 0.03 - 330 Hz) in a standard passive magnetically shielded room (AK3b, Vacuumschmelze, Germany) using a whole-head MEG (Elekta Neuromag Triux, Elekta Oy, Finland). Signals were captured by 102 magnetometers and 204 orthogonally placed planar gradiometers at 102 different positions.

MEG data were preprocessed and analyzed offline using Fieldtrip (Oostenveld et al., 2011), an open-source toolbox for Matlab (www.mathworks.com), and custom-written functions. Maxfilter (version 2.2.15, Elekta Neuromag, Finland) was applied to the continuous MEG raw data using a signal space separation algorithm (Taulu & Kajola, 2005; Taulu & Simola, 2006) provided by the MEG manufacturer. This allowed us to 1) automatically remove and interpolate the data of bad channels; 2) remove artifacts from the MEG signal (50 Hz line-noise and its harmonics, 16.7 Hz noise from a passing train, and muscle activity with origins beyond the head); and 3) compensate, offline, for changes in head position across blocks by realigning the data to a common standard head position (-trans default Maxfilter parameter) based on the head position measured at the beginning of each block. Thereafter, continuous data were visually inspected and sample points of extensive artifacts were stored separately for each block and participant.

Two different pipelines were used for further preprocessing of the MEG data, one optimized for applying independent component analysis (ICA), a second one for further data analysis. First, to optimize preprocessing of the data for ICA, MEG data were filtered offline using a 100 Hz lowpass filter (sinc FIR, kaiser window, order 3624, cutoff (−6 dB) 100 Hz) and a 1-Hz highpass filter (sinc FIR, kaiser window, order 908, cutoff (−6 dB) 1 Hz). After rejecting sample points covering extensive artifacts, runica ICA (the implementation from EEGLAB, Delorme and Makeig, 2004) was applied in order to identify components originating from eye movements and heartbeat. The second preprocessing pipeline was optimized for further data analysis. Specifically, cleaned MEG data were filtered offline using a 35 Hz lowpass filter (sinc FIR, kaiser window, order 3624, cutoff (−6 dB) 35 Hz). Data were downsampled to 256 Hz. Thereafter, a 0.1 Hz highpass filter was applied (sinc FIR, kaiser window, order 9056, cutoff (−6 dB) 0.1 Hz).

Filtered continuous data (originating from the second preprocessing pipeline) were cleaned from the ICs (on average two per participant). Data were segmented into 3-s epochs (−1.5 s to 1.5 s time-locked to the onset of the sound) and cleaned from artifacts that were identified via visual inspection (see above). For further analysis, we used deviant trials only. Trials in which deviants occurred on the left or right are, hereafter, referred to as Deviant Left or Deviant Right, respectively. Only those deviant trials for which participants responded once, correctly, and within the predefined response window (100 to 800 ms following the target), entered the analysis. On average across participants, >95% (SD ≤ 3.38) of the trials per condition were included in the analysis.

### 2.4 Data analyses

#### 2.4.1 Behavioral Analysis

The following trials were excluded from the analysis of behavioral data: trials without responses, trials including responses faster than 100 ms or slower than 800 ms after target onset, and trials that included more than one button press. On average across participants, >98% (SD ≤ 2.44) of the trials per condition were included in the analysis. Individual RTs and hit rates were calculated. Since individual reaction times typically show a skewed distribution, we used medians.

#### 2.4.2 Statistical analyses

We used jamovi (version 1.2.22.0, https://www.jamovi.org/) for the statistical analyses of behavioral data. To determine the distraction effect on RTs and hit rates, we applied two-way repeated-measures analyses of variance (ANOVAs) using the factors sound type (deviant, standard) and deviant location (left, right). Note that we used standards before the corresponding deviants for the analyses. To determine the behavioral effect of the spatial shift of attention on RTs and hit rates, we applied two-way repeated-measures ANOVAs using the factors congruence type (congruent, incongruent) and deviant side (left, right). Note that for statistical purposes, hit rates were transformed to rationalized arcsine units (rau) to account for non-normality of proportion data (Studebaker, 1985). We reported the generalized eta-squared (η^2^_G_; Bakeman, 2005) as a measure of effect size. Where appropriate, we reported mean values (M) and the standard error of the mean (SEM) in the format M ± SEM.

### 2.5 MEG Data Analyses

For the analyses of MEG data in sensor and source space, we solely used the signal recorded at the gradiometers. Data were detrended to remove slow frequency shifts.

#### 2.5.1 Sensor space analyses

##### 2.5.1.1 Time-frequency analysis

Time-frequency analysis was performed on the preprocessed single-trial data between 1 and 30 Hz (1 Hz steps) using Morlet wavelets with a width of 5 cycles, for each participant and condition, separately. The analysis time window was shifted in steps of 50 ms between -1.5 to 1.5 s relative to deviant onset. Note that power was first computed for each individual gradiometer. In a second step, the power for each gradiometer pair was summed in order to yield an orientation independent estimate. Thereafter, single-trial spectral power was log-transformed with a base of 10. For each participant, trials were averaged for Deviant Left and Deviant Right, separately.

In order to study the involuntary shift of spatial attention, we followed a methodological approach applied in research on voluntary spatial attention, in which alpha power was contrasted for the cued / attended target positions, namely attend left versus attend right (Banerjee et al., 2011; Deng et al., 2020; Haegens et al., 2011; Wöstmann et al., 2016). Accordingly, here we contrasted alpha power for the left and right attention-capturing, task-distracting deviants. In particular, we calculated an index of attentional modulation of alpha power - the Attentional Modulation Index (AMI; Wöstmann et al., 2016) for each time-frequency bin (−1.5 to 1.5 s; 5-30 Hz) and at each sensor: AMI = Deviant Left – Deviant Right. In General, the AMI reveals a spatially resolved measure of attention effects on alpha power at each sensor. A positive AMI indicates larger neural responses for Deviant Left trials. A negative AMI indicates larger responses for Deviant Right trials.

In order to statistically examine the difference in alpha power between the Deviant Left and Deviant Right conditions and to locate the effect, we performed non-parametric, cluster-based dependent-samples t-tests, with Monte-Carlo randomization (Maris & Oostenveld, 2007). The tests were restricted to the alpha frequency range (8 – 14 Hz) and the time window ranging from 0.2 to 0.6 s following the deviant sound and preceding the target (cf. Feng et al., 2017; Störmer et al., 2016; Weise et al., 2016). Since we had a priori hypotheses concerning the direction of the alpha power modulations expected in each hemisphere, we applied paired-sample, one-tailed Student’s t-tests for the left-hemispheric channels and the right-hemispheric channels, separately. The Student’s t-tests were applied for each sensor-time-frequency bin separately for each hemisphere by testing the alpha power values of Deviant Left against the alpha power values of Deviant Right. Based on this, clusters were formed among neighboring sensors of the respective hemisphere and corresponding time-frequency bins that exceeded an a priori chosen threshold (p = 0.05). The sum of the t-values within the respective cluster was used as the cluster-level statistic. By permuting the data between the two conditions and recalculating the test statistic 10,000 times, we obtained a reference distribution of maximum cluster t-values to evaluate the significance of the to-be-tested contrast. The power distributions between the two conditions were considered to be significantly different from each other if no more than 5% of the sum-t-values of all of the 10,000-permuted data were larger than the sum-t-value of the to-be-tested cluster. We applied Bonferroni correction to the significance thresholds as we used two statistical tests on non-independent data. Accordingly, the significance threshold was 0.025. Note that these tests were not performed on baseline-corrected data.

Moreover, we ran the time-frequency analysis a second time using the same parameters as described above, but with the preceding step that the evoked signal averaged across trials of the respective condition was subtracted in the time domain from every single trial, to increase sensitivity to non-phase locked signals, and to reduce contamination by phase-locked signals. The results are comparable to those obtained via the time-frequency analysis of phase-locked activity and can be found in the supplementary material (Figure S1).

Additionally, we calculated the laterality index (LI), i.e. the difference of the AMI (see above; Wöstmann et al., 2016) between the left and the right hemisphere, including either all channels of one hemisphere or only those of the region of interests (ROIs). Two ROIs - the parieto-occipital ROI and the centro-temporal ROI - had been identified via the preceding cluster-based permutation analysis and consisted of three channels that showed prominent deviant-related effects in the deviant–target interval in one hemisphere. To calculate the LI, we also selected the analogous three channels of the other hemisphere. In general the LI indicates a hemispheric lateralization of alpha power by the spatial attention demands under a given condition. To statistically test for that lateralization, we applied the Wilcoxon signed rank test to contrast the AMIs of the left hemispheric channels and the AMIs of the right hemispheric channels based on individual alpha power that was averaged across time, frequency and channels of the corresponding ROI or hemisphere (cf. Wöstman et al., 2016).

We further investigated whether the lateralized alpha power modulation observed in the data may be explained by a decrease or an increase with respect to a pre-stimulus baseline. Therefore, we analyzed the time course of alpha power separately for each condition (Deviant Left, Deviant Right) and region of interest (ROI; right-hemispheric centro-temporal ROI, left-hemispheric parieto-occipital ROI). The ROIs had been identified via the preceding cluster-based permutation analysis (see above). Each ROI (right-hemispheric centro-temporal ROI, left-hemispheric parieto-occipital ROI) consisted of three channels that showed prominent deviant-related effects in the deviant–target interval. Data were pooled within each ROI and across frequencies (8 – 14 Hz). Alpha power was submitted to non-parametric, cluster-based, dependent-samples t-tests with Monte-Carlo randomization (n = 10,000), contrasting the data at each sample point in the deviant-target interval (0.2 to 0.6 s following the deviant) with the mean alpha power of the baseline period (−0.4 to -0.2 s; Feng et al., 2017). Since we had no a priori hypotheses on the direction of the alpha power modulations expected in the respective ROI, we applied paired-sample, two-tailed Student’s t-tests for the cluster analysis (see above).

Note that for the analysis of alpha power we were exclusively interested in the activity following the deviant sound reflecting involuntary shifts of spatial attention towards the sound location. Accordingly, we did not expect differences between congruent and incongruent trials as the sounds were not predictive about the target location. Thus, there was no intent to contrast alpha activity between congruent and incongruent trials.

Also, note that the paradigm was not designed - and as a consequence is not suitable - to analyze alpha power between deviant and standard trials given that deviants were presented lateralized while standards were presented binaural.

#### 2.5.2 Source space analyses

##### 2.5.2.1 Source projection of time series data

For each participant, realistically shaped, single-shell head models (Nolte, 2003) were computed by co-registering the participants’ head shapes either with their structural MRI (18 participants) or—when no individual MRI was available (12 participants)—with a standard brain from the Montreal Neurological Institute (MNI, Montreal, Canada), warped to the individual head shape. A grid with 3-millimeter resolution based on an MNI template brain was morphed to fit the brain volume of each participant. The leadfield was calculated for each grid point and the forward model was computed for gradiometers only.

To project the preprocessed single-trial sensor data into source space (i.e., to the points of the grid), we applied the linearly constrained minimum variance (LCMV) spatial filter (Van Veen et al., 1997). We followed a procedure described for single virtual sensors (http://www.fieldtriptoolbox.org/tutorial/virtual_sensors/) and extended it to 146,400 points covering the gray matter of the brain. The covariance matrix across all trials (including Deviant Left and Deviant Right conditions) was calculated and used to obtain a LCMV beamformer spatial ?lter for each of the grid points (for a similar approach, see Neuling et al., 2015; Ruhnau et al., 2016). The covariance window for the calculation of the beamformer filter was based on a 2-s time window centered at deviant onset. To optimize the analysis in source space (i.e., increase spatial resolution using a high-definition grid and at the same time compensate for computation time), we divided the individual brain into 333 parcels (Gordon et al., 2016; http://www.nil.wustl.edu/labs/petersen/Resources.html) based on the ‘Anatomical Automatic Labeling’ [AAL] atlas (Tzourio-Mazoyer et al., 2002; provided by the fieldtrip toolbox). That is, for each anatomical area, the spatial filters of the corresponding grid points were averaged to obtain one spatial filter per parcel. The resulting ‘averaged filter’ was then multiplied with the sensor level data to obtain time-series data for each of the 333 parcels. A regularization parameter (lambda = 5%) was applied.

##### 2.5.2.3 Time-frequency analysis

For each participant, time-frequency analysis was performed on the single-trial data in source space between 1 and 30 Hz (1 Hz steps) using Morlet wavelets with a width of 5 cycles. The analysis time window was shifted in steps of 50 ms between -1.5 to 1.5 s relative to deviant onset. For each participant, trials were averaged separately for Deviant Left and Deviant Right.

To estimate the deviant-induced sensor level effects in source space, alpha power was averaged over frequency (8 -14 Hz) and time (0.2 - 0.6 s). T values were calculated for each of the parcels and the corresponding time-frequency bin, contrasting the alpha power values between Deviant Left and Deviant Right. Consistent with the original hypothesis and the sensor-space results, we only report positive T values in the right hemisphere and negative T values in the left hemisphere and applied a threshold corresponding to a one-sided student T test with p <= .05. Note that the T values calculated in source space serve only to visualize the effects already established in sensor space. They do not constitute statistical tests. T values were solely calculated because they are basically the difference between the two conditions normalized by the standard deviation, thereby controlling for different location dependent noise levels.

The respective labels of localized brain regions were identified with an anatomical brain atlas (AAL atlas; Tzourio-Mazoyer et al., 2002) and a network parcellation atlas (Gordon et al., 2016).

As with the sensor space analysis, we ran the time-frequency analysis and corresponding statistics a second time using the same parameters described above, but with the preceding step that the evoked signal averaged across trials of the respective condition was subtracted in the time domain from every single trial. The results are comparable to those obtained via the time-frequency analysis of phase-locked activity and can be found in the supplementary material (Figure S1).

## 3. Results

### 3.1 Behavioral data

In terms of accuracy, task performance was generally very good, as indicated by the high hit rate (>0.97) for targets, regardless of whether they were preceded by deviants or standards (Figure 2A). The two-way repeated measures ANOVA on hit rates shows a significant main effect for factor stimulus type (F(1, 29) = 18.51, p < .001, η^2^_G_ = .087), with higher hit rates for deviant trials (0.99 ± 0.003) than standard trials (0.97 ± 0.004). Neither was there a significant main effect for the factor deviant side (F(1, 29) = 1.09, p = .304, η^2^_G_ = .005), nor a significant interaction (F(1, 29) = .37, p = .548, η^2^_G_ = .001). This result may be linked to an increase in nonspecific arousal upon the occurrence of a deviant (Näätänen, 1992), which nevertheless facilitates task performance. Hit rates did not differ when deviant and target occurred on the congruent vs. incongruent side (Figure 2B), as indicated by the two-way repeated-measures ANOVA yielding neither significant main effects for the factors congruence type (F(1, 29) = 2.44, p = .129, η^2^_G_ = .017) and deviant side (F(1, 29) = .13, p = .726, η^2^_G_ = .001), nor a significant interaction (F(1, 29) = .06, p = .815, η^2^_G_ < .001).

**Figure 2:**
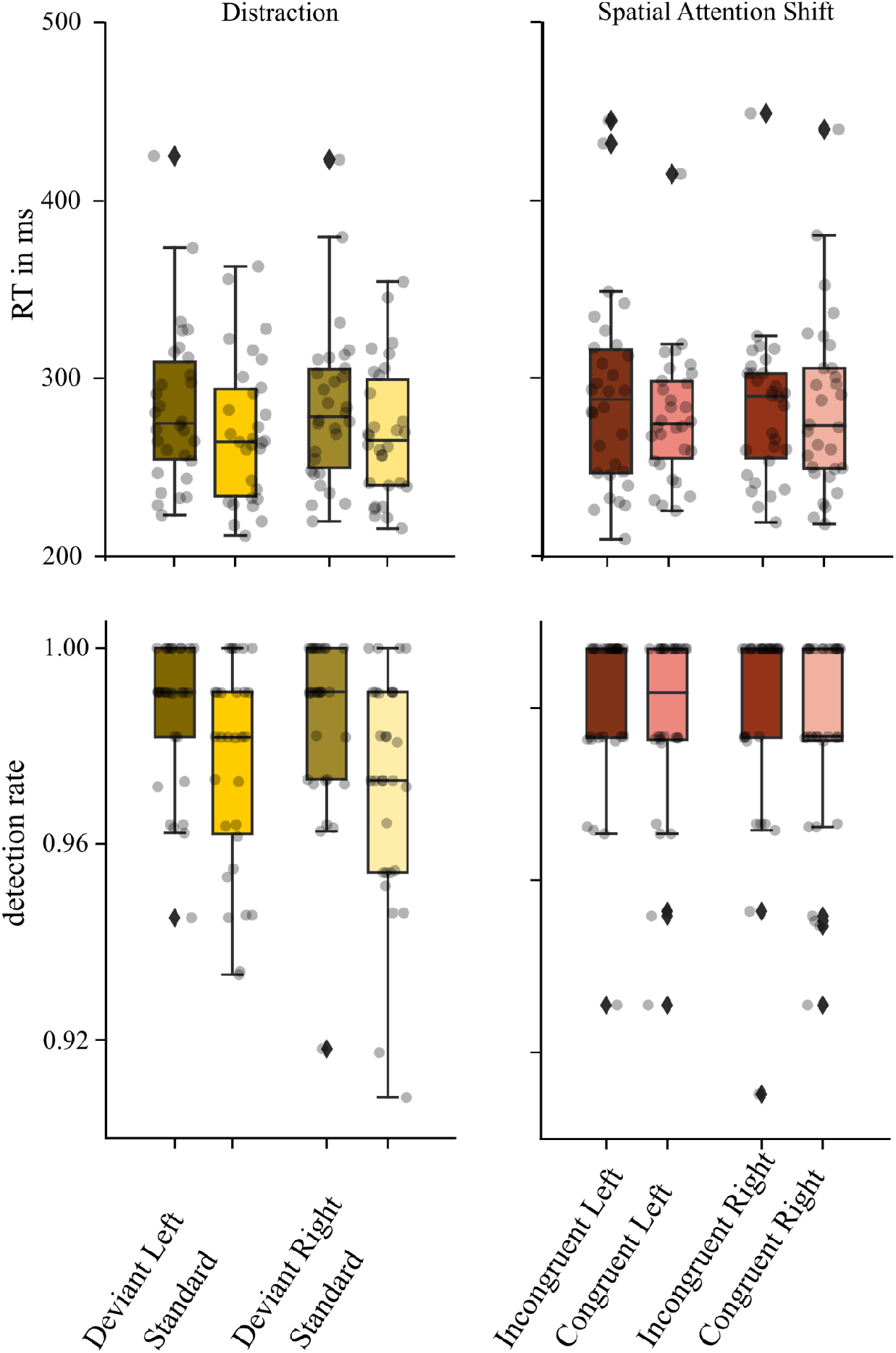
Boxplots of RTs and hit rates in response to visual targets to determine a distraction effect (left) or a shift of spatial attention (right) caused by the deviant. Gray dots represent individual data. (Left) RTs and hit rates for targets following deviants presented on the left or right side versus the standard presented before the corresponding deviant. (Right) RTs and hit rate for targets following deviants on the same versus different side in the Incongruent and Congruent conditions. Data were analyzed with respect to the deviant side (left, right).

As depicted in Figure 2A, RTs were prolonged when targets followed a deviant versus a standard. This is substantiated by the two-way repeated measures ANOVA on RTs, yielding a significant main effect for stimulus type (F(1, 29) = 24.62, p < .001, η^2^_G_ = .028; deviants: 296 ± 9 ms; standards: 278 ± 8 ms). There was neither a significant main effect for the factor deviant side (F(1, 29) = .34, p = .567, η^2^_G_ < .001), nor a significant interaction (F(1, 29) = 0.25, p = .624, η^2^_G_ < .001). Overall, these results point to a distraction effect due to an attention shift towards the task-distracting deviant. As depicted in Figure 2B, RTs were shorter when deviant and target occurred on the congruent side compared to the incongruent. This is substantiated by the two-way repeated measures ANOVA on RTs, yielding a significant main effect for congruence type (F(1, 29) = 4.28, p = .048, η^2^_G_ = .00; congruent: 293 ± 9 ms; incongruent: 298 ± 10 ms). There was neither a significant main effect for the factor deviant side F(1, 29) = .10, p = .757, η^2^_G_ < .001), nor a significant interaction (F(1, 29) = 1.06, p = .312, η^2^_G_ = .004). Overall, these results point to a *spatial* shift of attention towards the task-distracting deviant sound.

### 3.2 Alpha power analysis

We analyzed alpha-band (8 - 14 Hz) power to probe whether attentional reorienting towards the location of the deviant sound is also reflected in neural oscillatory activity, as indexed via the current RT data. We followed a methodological approach used to measure voluntary attention in which alpha power for the cued / attended target positions had been contrasted (attend left versus attend right; Banerjee et al., 2011; Deng et al., 2020; Haegens et al., 2011; Wöstmann et al., 2016). Accordingly, we contrasted alpha power for the left and right attention-capturing deviants (AMI, as defined earlier). A positive AMI indicates higher alpha power for Deviant Left trials and a negative AMI indicates higher alpha power for Deviant Right trials. We tested the effect of conditions (Deviant Left vs. Deviant Right) on the topographies of alpha power via non-parametric, cluster-based, permutation statistics. Since we had a priori hypotheses concerning the direction of the alpha power modulations expected in each hemisphere, we used the corresponding paired-sample, one-tailed Student’s t-tests on the left-hemispheric and the right-hemispheric channels, separately.

Overall, the results clearly show that alpha power to lateralized deviants is modulated in a spatially selective manner (Figure 3). In particular, the AMI was positive at channels over the left hemisphere (mean = .01, SD = .03) and negative at channels over the right hemisphere (mean = -.01, SD = .03). The results for the right hemispheric channels show that alpha power is significantly higher for Deviant Left trials compared to Deviant Right trials in the time window from 0.2 s to 0.6 s following the auditory deviant and preceding the visual target (see Figure 3 A). This effect is most pronounced over left parieto-occipital sensors (p = .023), as highlighted via the scalp distribution of the alpha power modulation (see Figure 3 C). The main generators of this sensor level effect are estimated in parieto-occipital regions (see Figure 3 D).

**Figure 3:**
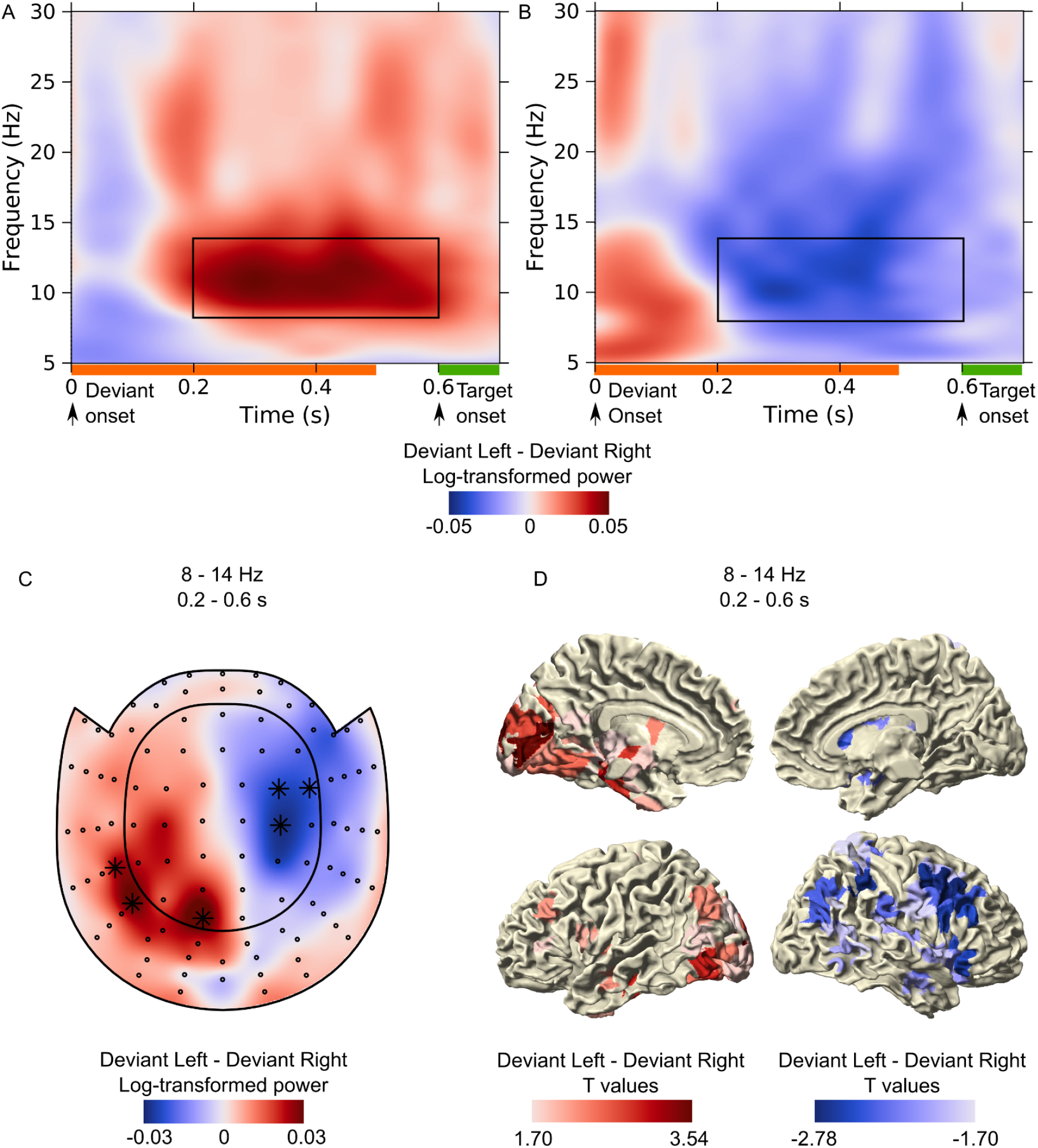
Log10-transformed alpha power modulation to lateralized deviants based on group-averaged data for Deviant Left minus Deviant Right. (A and B) Time-frequency representations highlighting (A) high alpha power ipsilateral to the deviant location at left-hemispheric posterior gradiometers (average across three channels showing a prominent effect, marked in C with a star), and (B) low alpha power contralateral to the deviant location at right-hemispheric centro-temporal gradiometer (average across three channels showing a prominent effect, marked in C with a star). The black box marks the time-frequency range (time: 0.2 - 0.6 s, frequency: 8 - 14 Hz) used for the statistical cluster analysis. (C) Topoplot of the alpha power distribution in the 0.2 - 0.6 s time window following deviant onset. Stars indicate channels on which there was a prominent statistical effect. (D) Source estimation of the alpha power effects. Note that the data were not baseline-corrected.

The results for the right hemispheric channels show that alpha power is significantly lower for Deviant Left trials compared to Deviant Right trials in the time window 0.2 s to 0.6 s following the auditory deviant and preceding the visual target (see Figure 3 B). This effect is most pronounced over right centro-temporal sensors (p = .012), as highlighted via the scalp distribution of the alpha power modulation (see Figure 3 C). The main generators of the sensor level effect are estimated in fronto-parietal regions (see Figure 3 D). Note that our analysis of non-phase-locked power yielded comparable results (Figure S1).

Further, we calculated the laterality index (LI) to test for a hemispheric lateralization of alpha power by the spatial attention demands under a given condition. Crucially, the LI, i.e. the difference of the left-hemispheric AMI and the right-hemispheric AMI, was significant (Wilcoxon signed-rank test; complete hemisphere: z = 3.66; p <.001; left-hemispheric posterior ROI: z = 4.24; p <.001; right-hemispheric centro-temporal ROI: z =3.83; p <.001).

To test whether the observed alpha lateralization (see Figure 3) was driven by a decrease or increase in alpha-band power relative to a pre-stimulus baseline, we analyzed the time courses of alpha power for the right hemispheric centro-temporal ROI and the left hemispheric parieto-occipital ROI separately for the left and right deviant. We contrasted the alpha power following the deviant with the mean alpha power of the baseline period, via a non-parametric cluster-based permutation statistic. Since we had no a priori hypotheses on the direction of the alpha power modulations, we applied paired-sample, two-tailed Student’s t-tests. The results show that the right deviant caused a significant increase in alpha power in the right-hemispheric centro-temporal ROI, in the time range 0.2 to 0.4 s after sound onset (p = .017, see Figure 4 A). Furthermore, for the left deviant, we found a significant increase in alpha power in the left-hemispheric parieto-occipital ROI, in the time range 0.2 to 0.3 s after sound onset (p = .039, see Figure 4 B). Additionally, the right deviant caused a significant decrease in alpha power in the left-hemispheric parieto-occipital ROI, in the time range 0.5 to 0.6 s after sound onset (p = .044, see Figure 4 B). The left deviant neither caused a significant increase nor decrease in alpha power in the left-hemispheric posterior ROI.

**Figure 4:**
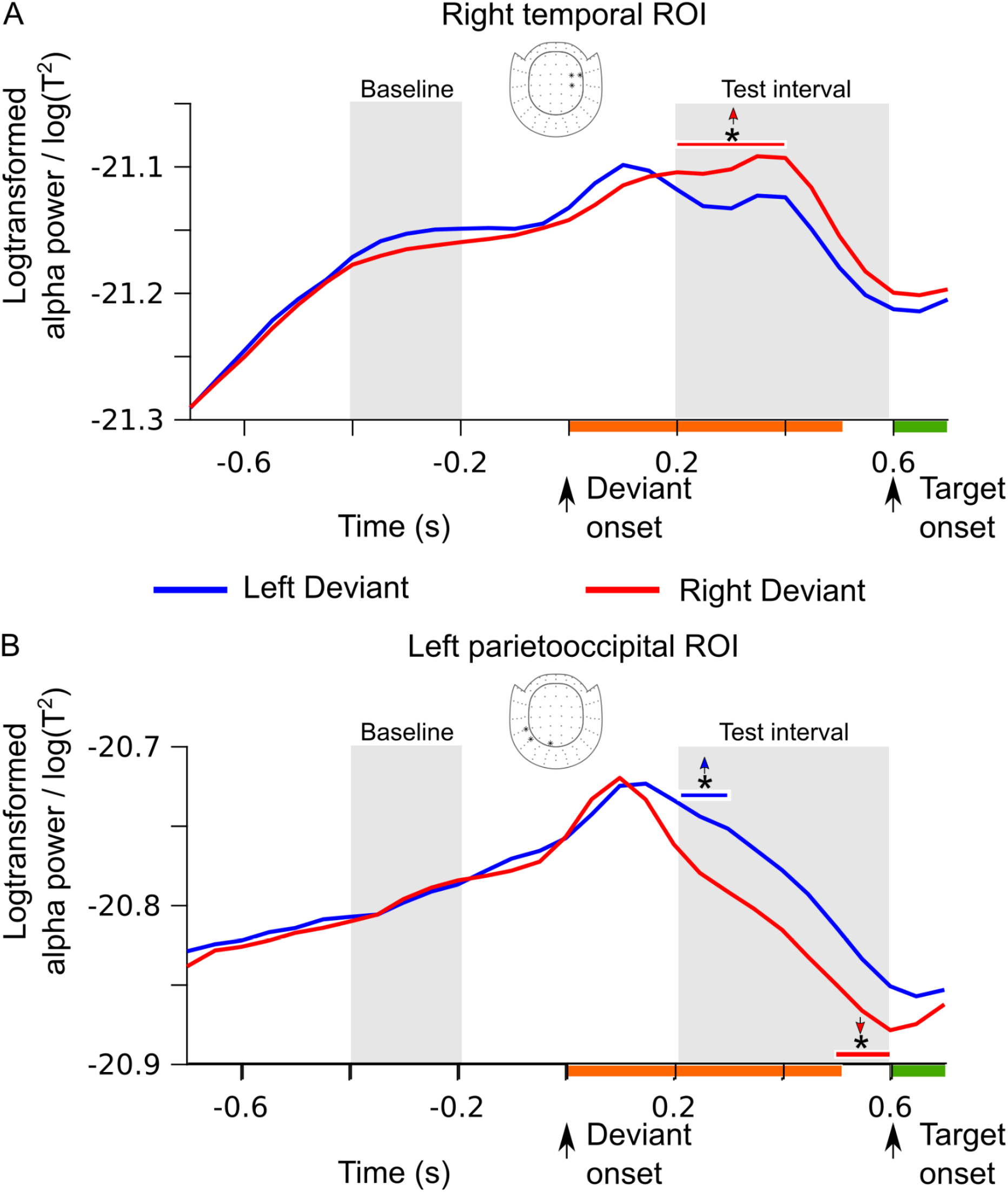
Time courses of log10-transformed and group-averaged alpha power for (A) the right-hemispheric centro-temporal ROI and (B) the left-hemispheric parieto-occipital ROI, time-locked to the left deviant (red line) or the right deviant (blue line). Channels belonging to the ROI are marked with stars in the corresponding channel layout. Significant time samples are illustrated by the horizontal red and blue lines (*p < .05). Upward directed arrows indicate an alpha power increase relative to baseline, and the downward directed arrow indicates an alpha power decrease with respect to baseline. Gray areas indicate the baseline interval (−0.4 to -0.2 s) and the test interval (0.2 to 0.6 s). The orange and green bars indicate the presentation time of deviant and target, respectively.

## 4. Discussion

Our behavioral reaction time and magnetoencephalographic alpha power data support the notion that task-distracting deviant sounds cause a shift in spatial attention that contributes to distraction.

### 4.1 The shift of involuntary *spatial* attention contributes to distraction

As expected, our distraction effect - that is, the longer RTs to targets that followed a deviant versus a standard - shows that the deviant captured attention involuntarily. This is in line with numerous findings that were obtained via a deviant distraction task (Bendixen et al., 2007; Parmentier et al., 2008; Schröger & Wolff, 1998; Wetzel et al., 2012).

Importantly, we observed an advantage in response speed when targets followed deviants on the same versus opposite side. This clearly speaks in favor of a *spatial* shift of attention, which has been proposed, though not directly tested, earlier (Parmentier et al., 2008). That is, when the deviant and target occur on the same side, the time penalty due to the spatial shift is lower compared to when both events occur on opposite sides. This finding is in line with behavioral data obtained via the stimulus-driven spatial cueing paradigm (McDonald et al., 2000), and receives further support from our MEG data, as will be discussed below.

Note that the spatial component of the attention shift unlikely accounts for the overall distraction effect in full; in fact, several cognitive determinants may contribute to it. A second determinant that could be at play in the current crossmodal situation, is the involuntary shift of attention across sensory modalities (i.e., from vision to audition). In this case, the time penalty accumulated by a shift of attention from the visual to the auditory modality (i.e., upon the onset of the deviant), and its re-orientation towards the visual modality (i.e., upon the onset of the target), would contribute to distraction (Parmentier, 2014). This idea receives support from behavioral studies showing that RTs are slower for a target in an unexpected modality compared to a target in an expected modality (Boulter, 1977; Spence et al., 2001).

At first glance, the RT data seem to be at odds with the hit rates. While the RTs indicate a decrease in performance for targets following a deviant versus a standard, the hit rates rather reflect an increase. However, this apparent contradiction can be reconciled by linking the RTs to the costs of the attention shift towards the deviant (Schröger & Wolff, 1998; Parmentier et al., 2008; for a review, see Parmentier 2014), and by linking the hit rates to an increase in nonspecific arousal that improves cognitive functioning (Näätänen, 1992). The current findings are well in line with previous research on deviant distraction, showing both costs and benefits (SanMiguel et al., 2010; Wetzel et al., 2012, Wetzel et al, 2019).

### 4.2 Posterior alpha power modulation reflects an involuntary shift of spatial attention

Our study is the first to show that a lateralized deviant sound modulates posterior alpha power in a spatially selective way. As expected, we observed a lateralized alpha power modulation over parieto-occipital sensors in the time window following the deviant sound and preceding the visual target. In line with our hypothesis, posterior alpha power modulation was higher in the hemisphere ipsilateral versus contralateral to the deviant location and likely reflects sound-induced spatial attention bias (involuntary spatial attention: Feng et al., 2017; Störmer et al., 2016; voluntary spatial attention: Deng et al., 2019; Deng et al., 2020; Frey et al., 2014; Wöstmann et al., 2016). This alpha power effect occurred relatively early and was rather short-lived, thereby closely matching the characteristics of involuntary attention both in timing and in the build-up phase (Corbetta & Shulman, 2002). The whole brain analyses further confirm the parieto-occipital localization of the effect (Figure 3 D).

Our data also show commonalities with recent work from the voluntary attention field in which auditory distractors were presented lateralized in space while the targets were presented centrally in front of the participant (Wöstmann et al., 2019; for a review, see Schneider et al., 2021). Similar to us, Wöstmann and colleagues (2019) found that lateralized auditory distractors induced lateralized posterior alpha power modulation (see Rösner et al., 2020 for converging findings in the visual domain). It is worth noting that in that study, alpha power modulation was lower in the hemisphere ipsilateral to the auditory distractor and higher in the hemisphere contralateral to the auditory distractor. At first glance, this may appear at odds with our pattern of results, since we found the exact opposite. However, it is important to recall that the two studies differ in important ways. Indeed, Wöstmann and colleagues (2019) addressed the issue of voluntary attention while our study examined involuntary attention, involving different cognitive mechanisms and designs. For example, the design of Wöstmann and colleagues (2019) made use of distractors that were presented concurrently with the targets on every trial. In addition, the location of the distractor (left vs. right) was manipulated across blocks and was therefore predictable. In contrast, our study used task-distracting deviants that preceded targets and their location was unpredictable from trial to trial. These crucial aspects may explain why the earlier data reflect the suppression of a distractor (e.g., the background noise when following a conversation at a cocktail party) while our results speak more to the issue of distractor / deviant selection (e.g. the ambulance siren when driving a car).

From a functional point of view, our posterior alpha power increase is also an interesting finding. A lot of what we know about the functional role of oscillatory alpha activity in the context of spatial attention originates from voluntary attention research. Via endogenous spatial cueing tasks, attention was voluntarily guided to the location of an upcoming target (visual target: Kelly et al., 2006; Rihs et al., 2009; Worden et al., 2000; auditory target: Banerjee et al., 2011; ElShafei et al., 2018; Frey et al., 2014; Müller & Weisz, 2012; Wöstman et al., 2019). In those tasks, lateralized alpha power was larger over the ipsilateral than the contralateral hemisphere, relative to the attended target location. This alpha power modulation was driven by 1) an alpha power decrease in the contralateral hemisphere and/or an alpha power increase ipsilateral to the attended side (e.g., Haegens et al., 2011); 2) a bilateral alpha power decrease that was maximal over the hemisphere contralateral to the attended position (e.g., Thut et al., 2006); or 3) a bilateral alpha power increase that was maximal over the hemisphere ipsilateral to the attended side (e.g., Banerjee et al., 2011; Worden et al., 2000). The pattern that is observed, reflecting mechanisms of suppression or enhancement at play in attention biasing, probably relies on divergent task demands of the respective study (Kelly et al., 2006; Thut et al., 2006). In any case, based on those studies, it has been widely acknowledged that decreased alpha power reflects the processing of task-relevant information in the voluntarily attended space, and increased alpha power indicates the inhibition of processes associated with task irrelevant information in the unattended space (for a review, see Klimesch, 2012). In a similar vein, our left-hemispheric parieto-occipital alpha power increase due to the left deviant may reflect the suppression of irrelevant information in the unattended space, that is, where involuntary spatial attention was *not* captured. This functional inhibition of posterior brain areas may free up resources for brain areas contralateral to the deviant site that process relevant information (Jensen & Mazaheri, 2010; Meeuwissen et al., 2011), which are also consequently involved in attention reorienting towards the deviant location. This interpretation receives support from our behavioral data, suggesting that deviants capture spatial attention (Figure 2B). Furthermore, it is in line with findings suggesting that a sound can engage visual processing at the same location (Feng et al., 2014; Feng et al., 2017; McDonald et al., 2013; Störmer et al., 2016). Interestingly, earlier work focussing on involuntary spatial attention exclusively found posterior alpha power decreases (Störmer et al., 2016; Feng et al., 2017). In more detail, the attention-capturing sound induced a bilateral alpha power decrease that was more prominent in the hemisphere contralateral to the location of the sound. Based on these earlier data, the authors suggested that a shift in involuntary attention exclusively facilitates target processing on the cued side, whereas target processing on the opposite side is not negatively impacted (Feng et al., 2017; Störmer et al., 2016). This conclusion was substantiated by the characteristic pattern of alpha power modulation preceding correct and incorrect discriminations of valid and invalid targets (Feng et al., 2017). Our results - demonstrating an increase of parieto-occipital alpha power - challenge that conclusion, as well as the prevalent notion that the “lack of an early inhibitory influence may represent a fundamental difference between involuntary orienting and voluntarily directed attention” (Feng et al., 2017, page 326). The difference between the current and previous studies is that our’s did not only apply an exogenous spatial cueing task, but also a distraction task, thus, likely resulting in different task demands (Kelly et al., 2006; Thut et al., 2006).

Considering the crossmodal nature of our task involving attention-capturing deviant sounds and visual targets, one might argue that our posterior alpha power increase was rather linked to the attentional disengagement from vision. Indeed, the outcome from intermodal attention tasks supports this view (Fox et al., 1998; Fu et al., 2001). For example, it is known that directing attention to auditory (versus visual) features of upcoming audio-visual targets increased posterior alpha power in the pre-target interval (Fox et al., 1998). That finding was interpreted to reflect “a disengaged visual attentional system in preparation for anticipated auditory input that is attentionally more relevant.” We cannot rule out that alternative account. However, we consider the spatial shift of attention explanation more likely. This receives support from our reaction time data from the congruent and incongruent conditions.

### 4.3 Centro-temporal alpha power increase reflects disengagement of motor and auditory areas

In addition to the current evidence pointing to a spatial shift of attention (see above), we unexpectedly found a right-hemispheric alpha power modulation over centro-temporal sites (Figure 3). This right-hemispheric effect was driven by an increase in alpha power, relative to baseline, due to the right deviant (Figure 4 A). Supported by the source data, this may reflect the disengagement of motor areas, as briefly outlined below (Figure 3 D) and adds to the recent discussion that anatomically separate alpha oscillations may enable different functionalizations in attention tasks (for a review, see Schneider et al., 2021). However, we want to emphasize that our interpretation is tentative.

The disengagement of motor areas in the context of distraction due to unexpected deviant sounds is well known (Dutra et al, 2018; Vasilev et al., 2019; Wessel et al., 2016; for a review, see Wessel & Aron, 2017). Accordingly, deviants recruit a ‘stopping network’ including the prefrontal cortex/ pre-supplementary motor area and temporarily interrupt the current action. That network has been linked to motor slowing such as slowing of RTs. Its putative purpose is to put the task at hand on hold to free up processing resources and allow an attention shift towards the unexpected deviant. Our behavioral and MEG data nicely fit that view and stimulate further research concerning the neuronal mechanisms engaged. Previous research linked a power increase in the delta (2–4 Hz) and low-theta (5–8 Hz) frequency bands to successful stopping (Huster et al., 2013; for a review, see Wessel & Aron, 2017), while a modulation in posterior alpha activity has been linked to attentional processing (Wessel & Aron, 2014). Our findings tentatively suggest that central alpha power may possibly play a role in motor suppression. This would be in line with research showing an alpha power increase during voluntary suppression of actions (Hummel et al., 2002; Sauseng et al., 2013).

In any case, if central alpha power does indeed implement deviant-induced ‘motor-stopping’ one might wonder whether our behavioral data reflect this rather than the spatial attention effect. This is unlikely, however. Even though the deviant-induced motor interruption might in general contribute to longer reaction times (compared to standards), this should, however, not obscure our RT data supporting a spatial shift of attention. The motor interruption is induced by deviants irrespective of their location and becomes evident after about 150 ms (Wessel & Aron, 2013). Importantly, the effect of the spatial shift of attention exclusively depends on whether the target follows the deviant at the same or different side. Another question that might raise concerns is whether the observed central alpha power modulation can be explained by preparatory processes related to the task. We consider that far-fetched because: 1) Participants responded to the left and right target with both hands, and the button-response assignment was counterbalanced across participants; 2) The alpha power modulation occurred in the pre-target interval, thus before the participant knew which hand was required for the response.

### 4.4 Conclusion

Here, we show that lateralized deviant sounds cause a shift of involuntary spatial attention and contribute to distraction. This is evident in our reaction time data, which demonstrates that responses are faster when the target follows an auditory deviant on the same side compared to the opposite sides. Additionally, our MEG data clearly show that oscillatory alpha power is shaped in a spatially selective manner. Most importantly, attention-capturing, task-distracting deviant sounds induced posterior alpha power that was higher in the left vs. right hemisphere. These attention-related changes were driven by a left-hemispheric posterior alpha power increase due to the left deviant, suggesting disengagement of visual areas that process task-irrelevant information outside the locus of involuntary attention.

Together with the behavioral data, our oscillatory data strengthen the view that alpha power reflects a shift of involuntary spatial attention.

## Author Note

This work was supported by the Deutsche Forschungsgemeinschaft (DFG, WE 6085/1-1, WE 6085/2-1). Fabrice Parmentier is an Adjunct Senior Lecturer at the University of Western Australia, and was supported by a research grant awarded to Fabrice Parmentier by the Ministry of Science, Innovation and Universities (PID2020-114117GB-I00), the Spanish State Agency for Research (AEI) and the European Regional Development Fund (FEDER). We thank Jonas Heilig for his help in collecting and preprocessing the data. We also thank Fred Seifter for his technical support in the lab. Further, we thank Reshanne Reeder and Kimberly Sherer for proofreading the manuscript. All authors disclose no potential sources of conflict of interest (financial or otherwise), which may be perceived as influencing an author’s objectivity. For reprints: Annekathrin Weise.

## Data and code availability

The preprocessed MEG and behavioral data used to generate the figures and calculate the statistics are available at the OSF public repository (https://osf.io/qt95u/). The corresponding data analysis code is available at the corresponding author’s GitLab repository (https://gitlab.com/aweise/crossfrog-final-analysis). The original raw data are available, upon reasonable request, from the corresponding author.

## Supplementary materials

**Figure S1:**
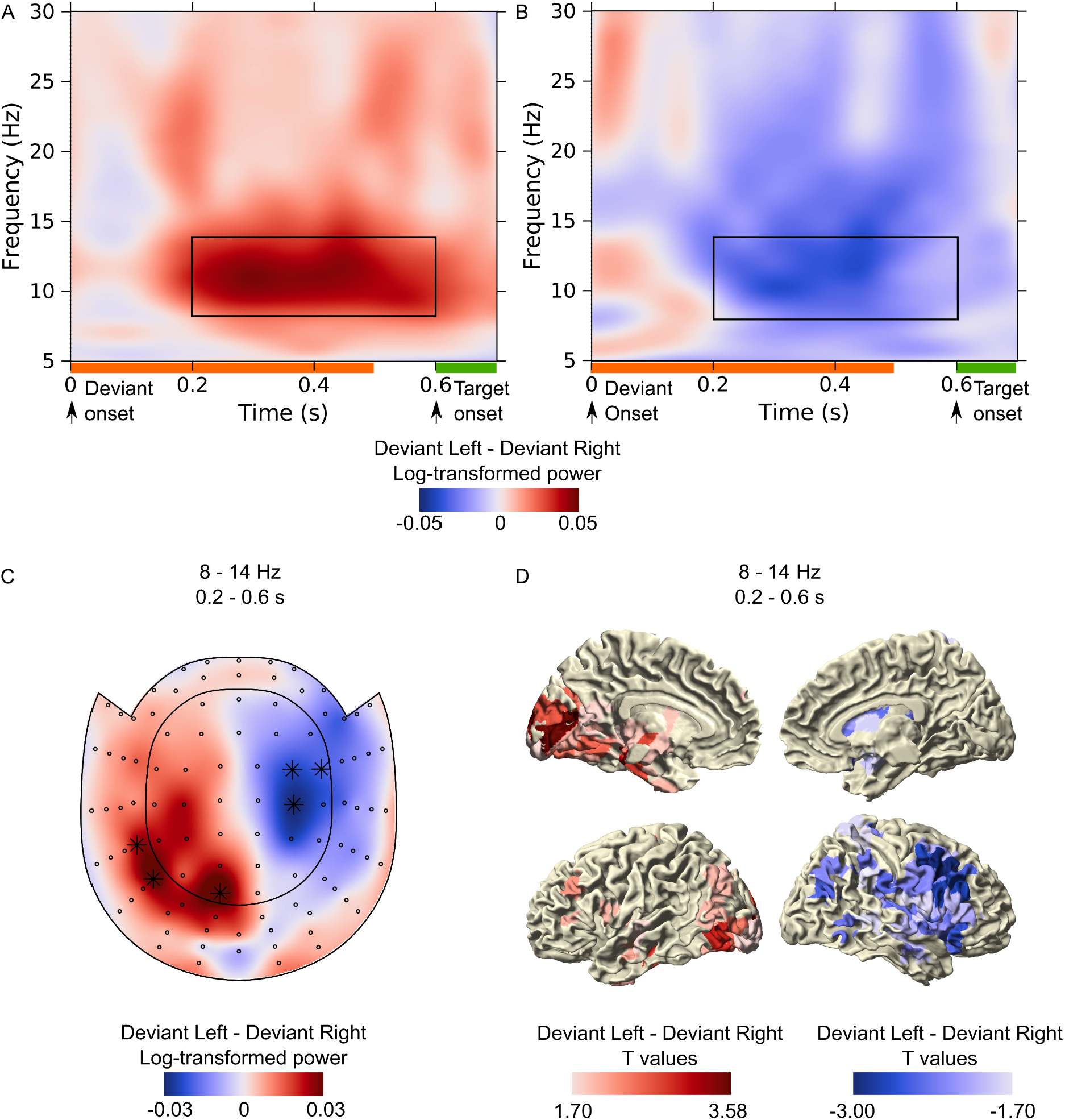
Log10-transformed, non-phase locked alpha power modulation to lateralized deviants based on group-averaged data for Deviant Left minus Deviant Right. (A and B) Time-frequency representations highlighting (A) high alpha power ipsilateral to the deviant location at left-hemispheric posterior gradiometers (average across three channels showing a prominent effect, marked in C with a star), and (B) low alpha power contralateral to the deviant location at right-hemispheric centro-temporal gradiometer (average across three channels showing a prominent effect, marked in C with a star). The black box marks the time-frequency range (time: 0.2 - 0.6 s, frequency: 8 - 14 Hz) used for the statistical cluster analysis. (C) Topoplot of the alpha power distribution in the 0.2 - 0.6 s time window following deviant onset. Stars indicate channels on which there was a prominent statistical effect. (D) Source estimation of the alpha power effects. Note that the data were not baseline-corrected.

## References

Bakeman, R. (2005). Recommended effect size statistics for repeated measures designs. Behavior Research Methods, 37(3), 379–384. https://www.doi.org/10.3758/bf03192707

Banerjee, S., Snyder, A. C., Molholm, S., & Foxe, J. J. (2011). Oscillatory alpha-band mechanisms and the deployment of spatial attention to anticipated auditory and visual target locations: Supramodal or sensory-specific control mechanisms? The Journal of Neuroscience, 31(27), 9923–9932. https://www.doi.org/10.1523/JNEUROSCI.4660-10.2011

Bendixen, A., Prinz, W., Horváth, J., Trujillo-Barreto, N. J., & Schröger, E. (2008). Rapid extraction of auditory feature contingencies. NeuroImage, 41(3), 1111–1119. https://www.doi.org/10.1016/j.neuroimage.2008.03.040

Bendixen, A., Roeber, U., & Schröger, E. (2007). Regularity extraction and application in dynamic auditory stimulus sequences. Journal of Cognitive Neuroscience, 19(10), 1664–1677. https://www.doi.org/10.1162/jocn.2007.19.10.1664

Boulter, L. R. (1977). Attention and reaction times to signals of uncertain modality. Journal of Experimental Psychology. Human Perception and Performance, 3(3), 379–388. https://www.doi.org/10.1037//0096-1523.3.3.379

Brainard, D. H. (1997). The Psychophysics Toolbox. Spatial Vision, 10(4), 433–436. https://www.doi.org/10.1163/156856897X00357

Cheal, M. L., & Gregory, M. (1997). Evidence of limited capacity and noise reduction with single-element displays in the location-cuing paradigm. Journal of Experimental Psychology. Human Perception and Performance, 23(1), 51–71. https://www.doi.org/10.1037//0096-1523.23.1.51

Corbetta, M., & Shulman, G. L. (2002). Control of goal-directed and stimulus-driven attention in the brain. Nature Reviews Neuroscience, 3(3), 201–215.

Corral, M.-J., & Escera, C. (2008). Effects of sound location on visual task performance and electrophysiological measures of distraction. Neuroreport, 19(15), 1535–1539. https://www.doi.org/10.1097/WNR.0b013e3283110416

Delorme, A., & Makeig, S. (2004). EEGLAB: An open source toolbox for analysis of single-trial EEG dynamics including independent component analysis. Journal of Neuroscience Methods, 134(1), 9–21. https://doi.org/10.1016/j.jneumeth.2003.10.009

Deng, Y., Choi, I., & Shinn-Cunningham, B. (2020). Topographic specificity of alpha power during auditory spatial attention. NeuroImage, 207, 116360. https://www.doi.org/10.1016/j.neuroimage.2019.116360

Deng, Y., Reinhart, R. M., Choi, I., & Shinn-Cunningham, B. G. (2019). Causal links between parietal alpha activity and spatial auditory attention. ELife, 8. https://www.doi.org/10.7554/eLife.51184

Dufour, A. (1999). Importance of attentional mechanisms in audiovisual links. Experimental Brain Research, 126(2), 215–222. https://www.doi.org/10.1007/s002210050731

Dutra, I. C., Waller, D. A., & Wessel, J. R. (2018). Perceptual surprise improves action stopping by nonselectively suppressing motor activity via a neural mechanism for motor inhibition. Journal of Neuroscience, 38(6), 1482–1492. https://www.doi.org/10.1523/JNEUROSCI.3091-17.2017

ElShafei, H. A., Bouet, R., Bertrand, O., & Bidet-Caulet, A. (2018). Two sides of the same coin: Distinct sub-bands in the alpha rhythm reflect facilitation and suppression mechanisms during auditory anticipatory attention. ENeuro, ENEURO.0141-18.2018. https://www.doi.org/10.1523/ENEURO.0141-18.2018

Escera, C., Alho, K., Schröger, E., & Winkler, I. (2000). Involuntary attention and distractibility as evaluated with event-related brain potentials. Audiology and Neuro-Otology, 5(3–4), 151–166. https://www.doi.org/10.1159/000013877

Escera, C., Alho, K., Winkler, I., & Näätänen, R. (1998). Neural mechanisms of involuntary attention to acoustic novelty and change. Journal of Cognitive Neuroscience, 10(5), 590–604. https://www.doi.org/10.1162/089892998562997

Feng, W., Störmer, V. S., Martinez, A., McDonald, J. J., & Hillyard, S. A. (2014). Sounds activate visual cortex and improve visual discrimination. The Journal of Neuroscience, 34(29), 9817–9824. https://www.doi.org/10.1523/JNEUROSCI.4869-13.2014

Feng, W., Störmer, V. S., Martinez, A., McDonald, J. J., & Hillyard, S. A. (2017). Involuntary orienting of attention to a sound desynchronizes the occipital alpha rhythm and improves visual perception. NeuroImage, 150, 318–328. https://www.doi.org/10.1016/j.neuroimage.2017.02.033

Foxe, J. J., Simpson, G. V., & Ahlfors, S. P. (1998). Parieto-occipital approximately 10 Hz activity reflects anticipatory state of visual attention mechanisms. Neuroreport, 9(17), 3929–3933.

Frey, J. N., Mainy, N., Lachaux, J.-P., Müller, N., Bertrand, O., & Weisz, N. (2014). Selective modulation of auditory cortical alpha activity in an audiovisual spatial attention task. The Journal of Neuroscience, 34(19), 6634–6639. https://www.doi.org/10.1523/JNEUROSCI.4813-13.2014

Fu, K.-M. G., Foxe, J. J., Murray, M. M., Higgins, B. A., Javitt, D. C., & Schroeder, C. E. (2001). Attention-dependent suppression of distracter visual input can be cross-modally cued as indexed by anticipatory parieto–occipital alpha-band oscillations. Cognitive Brain Research, 12(1), 145–152. https://www.doi.org/10.1016/S0926-6410(01)00034-9

Gordon, E. M., Laumann, T. O., Adeyemo, B., Huckins, J. F., Kelley, W. M., & Petersen, S. E. (2016). Generation and evaluation of a cortical area parcellation from resting-state correlations. Cerebral Cortex, 26(1), 288–303. https://www.doi.org/10.1093/cercor/bhu239

Haegens, S., Händel, B. F., & Jensen, O. (2011). Top-down controlled alpha band activity in somatosensory areas determines behavioral performance in a discrimination task. Journal of Neuroscience, 31(14), 5197–5204. https://www.doi.org/10.1523/JNEUROSCI.5199-10.2011

Hanslmayr, S., Gross, J., Klimesch, W., & Shapiro, K. L. (2011). The role of α oscillations in temporal attention. Brain Research Reviews, 67(1–2), 331–343. https://www.doi.org/10.1016/j.brainresrev.2011.04.002

Hartmann, T., & Weisz, N. (2020). An Introduction to the Objective Psychophysics Toolbox. Frontiers in Psychology, 11, 585437. https://www.doi.org/10.3389/fpsyg.2020.585437

Hummel, F., Andres, F., Altenmüller, E., Dichgans, J., & Gerloff, C. (2002). Inhibitory control of acquired motor programmes in the human brain. Brain: A Journal of Neurology, 125(Pt 2), 404–420. https://www.doi.org/10.1093/brain/awf030

Huster, R. J., Enriquez-Geppert, S., Lavallee, C. F., Falkenstein, M., & Herrmann, C. S. (2013). Electroencephalography of response inhibition tasks: Functional networks and cognitive contributions. International Journal of Psychophysiology: Official Journal of the International Organization of Psychophysiology, 87(3), 217–233. https://www.doi.org/10.1016/j.ijpsycho.2012.08.001

Jensen, O., & Mazaheri, A. (2010). Shaping functional architecture by oscillatory alpha activity: Gating by inhibition. Frontiers in Human Neuroscience, 4. https://www.doi.org/10.3389/fnhum.2010.00186

Kelly, S. P., Lalor, E. C., Reilly, R. B., & Foxe, J. J. (2006). Increases in alpha oscillatory power reflect an active retinotopic mechanism for distracter suppression during sustained visuospatial attention. Journal of Neurophysiology, 95(6), 3844–3851. https://www.doi.org/10.1152/jn.01234.2005

Kleiner, M., Brainard, D., & Pelli, D. (2007). “What’s new in Psychtoolbox-3?”. Perception, 36. ECVP Abstract Supplement. Abgerufen von ECVP Abstract Supplement.

Klimesch, W. (2012). Alpha-band oscillations, attention, and controlled access to stored information. Trends in Cognitive Sciences, 16(12), 606–617. https://www.doi.org/10.1016/j.tics.2012.10.007

Kontsevich, L. L., & Tyler, C. W. (1999). Bayesian adaptive estimation of psychometric slope and threshold. Vision Research, 39(16), 2729–2737. https://www.doi.org/10.1016/S0042-6989(98)00285-5

Luck, S. J., Hillyard, S. A., Mouloua, M., & Hawkins, H. L. (1996). Mechanisms of visual-spatial attention: Resource allocation or uncertainty reduction? Journal of Experimental Psychology. Human Perception and Performance, 22(3), 725–737. https://www.doi.org/10.1037//0096-1523.22.3.725

Maris, E., & Oostenveld, R. (2007). Nonparametric statistical testing of EEG-and MEG-data. Journal of Neuroscience Methods, 164(1), 177–190. https://www.doi.org/10.1016/j.jneumeth.2007.03.024

McDonald, J. J., Teder-Sälejärvi, W. A., & Hillyard, S. A. (2000). Involuntary orienting to sound improves visual perception. Nature, 407(6806), 906–908. https://www.doi.org/10.1038/35038085

McDonald, John J., Störmer, V. S., Martinez, A., Feng, W., & Hillyard, S. A. (2013). Salient sounds activate human visual cortex automatically. The Journal of Neuroscience, 33(21), 9194–9201. https://www.doi.org/10.1523/JNEUROSCI.5902-12.2013

Meeuwissen, E. B., Takashima, A., Fernández, G., & Jensen, O. (2011). Increase in posterior alpha activity during rehearsal predicts successful long-term memory formation of word sequences. Human Brain Mapping, 32(12), 2045–2053. https://www.doi.org/10.1002/hbm.21167

Müller, N., & Weisz, N. (2012). Lateralized auditory cortical alpha band activity and interregional connectivity pattern reflect anticipation of target sounds. Cerebral Cortex, 22(7), 1604–1613. https://www.doi.org/10.1093/cercor/bhr232

Näätänen, R. (1992). Attention and brain function. Hillsdale, NJ: Erlbaum.

Neuling, T., Ruhnau, P., Fuscà, M., Demarchi, G., Herrmann, C. S., & Weisz, N. (2015). Friends, not foes: Magnetoencephalography as a tool to uncover brain dynamics during transcranial alternating current stimulation. NeuroImage, 118, 406–413. https://www.doi.org/10.1016/j.neuroimage.2015.06.026

Nolte, G. (2003). The magnetic lead field theorem in the quasi-static approximation and its use for magnetoencephalography forward calculation in realistic volume conductors. Physics in Medicine and Biology, 48(22), 3637. https://www.doi.org/10.1088/0031-9155/48/22/002

Parmentier, F. B. R. (2014). The cognitive determinants of behavioral distraction by deviant auditory stimuli: A review. Psychological Research, 78(3), 321–338. https://www.doi.org/10.1007/s00426-013-0534-4

Parmentier, F. B. R., Elford, G., Escera, C., Andrés, P., & San Miguel, I. (2008). The cognitive locus of distraction by acoustic novelty in the cross-modal oddball task. Cognition, 106(1), 408–432. https://www.doi.org/10.1016/j.cognition.2007.03.008

Parmentier, F. B. R., Elsley, J. V., Andrés, P., & Barceló, F. (2011). Why are auditory novels distracting? Contrasting the roles of novelty, violation of expectation and stimulus change. Cognition, 119(3), 374–380. https://www.doi.org/10.1016/j.cognition.2011.02.001

Peylo, C., Hilla, Y., & Sauseng, P. (in press). Cause or consequence? Alpha oscillations in visuospatial attention. Trends in Neurosciences. https://www.doi.org/10.1016/j.tins.2021.05.004

Rihs, T. A., Michel, C. M., & Thut, G. (2009). A bias for posterior alpha-band power suppression versus enhancement during shifting versus maintenance of spatial attention. NeuroImage, 44(1), 190–199. https://www.doi.org/10.1016/j.neuroimage.2008.08.022

Rösner, M., Arnau, S., Skiba, I., Wascher, E., & Schneider, D. (2020). The spatial orienting of the focus of attention in working memory makes use of inhibition: Evidence by hemispheric asymmetries in posterior alpha oscillations. Neuropsychologia, 142, 107442. https://doi.org/10.1016/j.neuropsychologia.2020.107442

Ruhnau, P., Herrmann, B., Maess, B., Brauer, J., Friederici, A. D., & Schröger, E. (2013). Processing of complex distracting sounds in school-aged children and adults: Evidence from EEG and MEG data. Frontiers in Psychology, 4. https://www.doi.org/10.3389/fpsyg.2013.00717

Ruhnau, P., Neuling, T., Fuscá, M., Herrmann, C. S., Demarchi, G., & Weisz, N. (2016). Eyes wide shut: Transcranial alternating current stimulation drives alpha rhythm in a state dependent manner. Scientific Reports, 6, 27138. https://www.doi.org/10.1038/srep27138

Sanchez, G., Lecaignard, F., Otman, A., Maby, E., & Mattout, J. (2016). Active SAmpling Protocol (ASAP) to Optimize Individual Neurocognitive Hypothesis Testing: A BCI-Inspired Dynamic Experimental Design. Frontiers in Human Neuroscience, 10. https://www.doi.org/10.3389/fnhum.2016.00347

SanMiguel, I., Linden, D., & Escera, C. (2010). Attention capture by novel sounds: Distraction versus facilitation. European Journal of Cognitive Psychology, 22(4), 481–515. https://www.doi.org/10.1080/09541440902930994

Sauseng, P., Klimesch, W., Stadler, W., Schabus, M., Doppelmayr, M., Hanslmayr, S., … Birbaumer, N. (2005). A shift of visual spatial attention is selectively associated with human EEG alpha activity. European Journal of Neuroscience, 22(11), 2917–2926. https://www.doi.org/10.1111/j.1460-9568.2005.04482.x

Sauseng, Paul, Gerloff, C., & Hummel, F. C. (2013). Two brakes are better than one: The neural bases of inhibitory control of motor memory traces. NeuroImage, 65, 52–58. https://www.doi.org/10.1016/j.neuroimage.2012.09.048

Schneider, D., Herbst, S. K., Klatt, L.-I., & Wöstmann, M. (2021). Target enhancement or distractor suppression? Functionally distinct alpha oscillations form the basis of attention. European Journal of Neuroscience, 00, 1–10. https://doi.org/10.1111/ejn.15309

Schröger, E. (1996). A neural mechanism for involuntary attention shifts to changes in auditory stimulation. Journal of Cognitive Neuroscience, 8, 527–539. https://www.doi.org/10.1162/jocn.1996.8.6.527

Schröger, E., & Wolff, C. (1998). Behavioral and electrophysiological effects of task-irrelevant sound change: A new distraction paradigm. Brain Research. Cognitive Brain Research, 7(1), 71–87.

Shiu, L., & Pashler, H. (1994). Negligible effect of spatial precuing on identification of single digits. Journal of Experimental Psychology: Human Perception and Performance, 20(5), 1037–1054. https://www.doi.org/10.1037/0096-1523.20.5.1037

Spence, C., & Driver, J. (1997). Audiovisual links in exogenous covert spatial orienting. Perception & Psychophysics, 59(1), 1–22. https://www.doi.org/10.3758/BF03206843

Spence, C., Nicholls, M. E. R., & Driver, J. (2001). The cost of expecting events in the wrong sensory modality. Perception & Psychophysics, 63(2), 330–336. https://www.doi.org/10.3758/BF03194473

Störmer, V., Feng, W., Martinez, A., McDonald, J., & Hillyard, S. (2016). Salient, irrelevant sounds reflexively induce alpha rhythm desynchronization in parallel with slow potential shifts in visual cortex. Journal of Cognitive Neuroscience, 28(3), 433–445. https://www.doi.org/10.1162/jocn_a_00915

Studebaker, G. A. (1985). A “rationalized” arcsine transform. Journal of Speech and Hearing Research, 28(3), 455–462. https://www.doi.org/10.1044/jshr.2803.455

Taulu, S., & Simola, J. (2006). Spatiotemporal signal space separation method for rejecting nearby interference in MEG measurements. Physics in Medicine and Biology, 51(7), 1759–1768. https://www.doi.org/10.1088/0031-9155/51/7/008

Taulu, Samu, & Kajola, M. (2005). Presentation of electromagnetic multichannel data: The signal space separation method. Journal of Applied Physics, 97(12), 124905. https://www.doi.org/10.1063/1.1935742

Thut, G., Nietzel, A., Brandt, S. A., & Pascual-Leone, A. (2006). Alpha-band electroencephalographic activity over occipital cortex indexes visuospatial attention bias and predicts visual target detection. Journal of Neuroscience, 26(37), 9494–9502. https://www.doi.org/10.1523/JNEUROSCI.0875-06.2006

Tzourio-Mazoyer, N., Landeau, B., Papathanassiou, D., Crivello, F., Etard, O., Delcroix, N., … Joliot, M. (2002). Automated anatomical labeling of activations in SPM using a macroscopic anatomical parcellation of the MNI MRI single-subject brain. NeuroImage, 15(1), 273–289. https://www.doi.org/10.1006/nimg.2001.0978

Van Veen, B. D., van Drongelen, W., Yuchtman, M., & Suzuki, A. (1997). Localization of brain electrical activity via linearly constrained minimum variance spatial filtering. IEEE Transactions on Bio-Medical Engineering, 44(9), 867–880. https://www.doi.org/10.1109/10.623056

Vasilev, M. R., Parmentier, F. B., Angele, B., & Kirkby, J. A. (2019). Distraction by deviant sounds during reading: An eye-movement study. Quarterly Journal of Experimental Psychology (2006), 72(7), 1863–1875. https://www.doi.org/10.1177/1747021818820816

Weise, A., Hartmann, T., Schröger, E., Weisz, N., & Ruhnau, P. (2016). Cross-modal distractors modulate oscillatory alpha power: The neural basis of impaired task performance. Psychophysiology, 53(11), 1651–1659. https://www.doi.org/10.1111/psyp.12733

Wessel, J. R., & Aron, A. R. (2013). Unexpected Events Induce Motor Slowing via a Brain Mechanism for Action-Stopping with Global Suppressive Effects. Journal of Neuroscience, 33(47), 18481–18491. https://www.doi.org/10.1523/JNEUROSCI.3456-13.2013

Wessel, J. R., & Aron, A. R. (2017). On the globality of motor suppression: Unexpected events and their influence on behavior and cognition. Neuron, 93(2), 259–280. https://www.doi.org/10.1016/j.neuron.2016.12.013

Wessel, J. R., Jenkinson, N., Brittain, J.-S., Voets, S. H. E. M., Aziz, T. Z., & Aron, A. R. (2016). Surprise disrupts cognition via a fronto-basal ganglia suppressive mechanism. Nature Communications, 7, 11195. https://www.doi.org/10.1038/ncomms11195

Wetzel, N., Scharf, F., & Widmann, A. (2019). Can’t Ignore—Distraction by task-irrelevant sounds in early and middle childhood. Child Development, 90(6), e819–e830. https://www.doi.org/10.1111/cdev.13109

Wetzel, N., Widmann, A., & Schröger, E. (2012). Distraction and facilitation—Two faces of the same coin? Journal of Experimental Psychology. Human Perception and Performance, 38(3), 664–674. https://www.doi.org/10.1037/a0025856

Worden, M. S., Foxe, J. J., Wang, N., & Simpson, G. V. (2000). Anticipatory biasing of visuospatial attention indexed by retinotopically specific alpha-band electroencephalography increases over occipital cortex. The Journal of Neuroscience, 20(6), RC63.

Wöstmann, M., Alavash, M., & Obleser, J. (2019). Alpha oscillations in the human brain implement distractor suppression independent of target selection. Journal of Neuroscience, 39(49), 9797–9805. https://doi.org/10.1523/JNEUROSCI.1954-19.2019

Wöstmann, M., Herrmann, B., Maess, B., & Obleser, J. (2016). Spatiotemporal dynamics of auditory attention synchronize with speech. Proceedings of the National Academy of Sciences, 113(14), 3873–3878. https://www.doi.org/10.1073/pnas.1523357113

